# Comparative genomics reveals signature regions used to develop a robust and sensitive multiplex TaqMan real-time qPCR assay to detect the genus *Dickeya* and *Dickeya dianthicola*

**DOI:** 10.1101/847590

**Authors:** Shefali Dobhal, Gamze Boluk, Brooke Babler, Michael J. Stulberg, John Rascoe, Mark Nakhla, Toni A. Chapman, Alex B. Crockford, Michael Melzer, Anne M. Alvarez, Mohammad Arif

## Abstract

**Aims:** *Dickeya* species are high consequence plant pathogenic bacteria listed among the quarantine pathogens of the European Union; associated with potato disease outbreaks and subsequent economic losses worldwide. Early, accurate, and reliable detection of *Dickeya* spp. is needed to prevent establishment and further dissemination of this pathogen. Therefore, a multiplex TaqMan qPCR was developed for sensitive detection of *Dickeya* spp. and specifically, *D. dianthicola*.

**Methods and Results:** A signature genomic region for the genus *Dickeya* (*mglA/mglC*) and unique genomic region for *D. dianthicola* (alcohol dehydrogenase) were identified using a whole genome based comparative genomics approach. The developed multiplex TaqMan qPCR was validated using extensive inclusivity and exclusivity panels, and naturally/artificially infected samples to confirm broad range detection capability and specificity. Both sensitivity and spiked assays showed detection limit of 10 fg DNA.

**Conclusion:** The developed multiplex assay is sensitive and reliable to detect *Dickeya* spp. and *D. dianthicola* with no false positives or false negatives. It was able to detect mixed infection from naturally and artificially infected plant materials.

**Significance and Impact:** The developed assay will serve as a practical tool for screening of propagative material, monitoring the presence and distribution, and quantification of target pathogens in a breeding program. The assay also has applications in routine diagnostics, biosecurity and microbial forensics.

## Introduction

The genus *Dickeya* belongs to the family Enterobacteriaceae and is comprised of pectinolytic Gram negative plant pathogenic bacteria, responsible for the soft rot diseases worldwide. Due to their association with catastrophic diseases, *Dickeya* species have been listed among the top ten most destructive bacterial plant pathogens (Mansfield *et al*. 2012). Soft rot Enterobacteria are also associated with the most serious problems facing potato production worldwide. The impact of soft rot disease varies from country to country; in the Netherlands, the annual estimated losses are around €30 million per year, while in Israel, potato yield losses of 20-25% have been observed. (Tsror (Lahkim) *et al*. 2009; Raoul des Essarts *et al*. 2016; Buttimer *et al*. 2017).

The genus *Dickeya* originally included six species: *D. chrysanthemi, D. dadantii, D. dieffenbachiae, D. dianthicola, D. paradisiaca*, and *D. zeae* (Samson *et al*. 2005). Later, *D. dieffenbachiae* was reclassified as a subspecies of *D. dadantii* based on the DNA-DNA hybridization and phylogenetic analyses (Brady *et al*. 2012). *Dickeya* now has eight species including, *D. solani* (Van der Wolf *et al*. 2014), *D. aquatica* (Parkinson *et al*. 2014) and the recently described *D. fangzondai* (*Tian et al*. 2016). *Dickeya* species have a broad host range, infecting both monocots and dicots (Ma *et al*. 2007). They are responsible for soft rots of vegetables and ornamentals, resulting in high economic losses in various geographical locations worldwide (Van Vaerenbergh *et al*. 2012).

Symptoms caused by *Dickeya* spp. are more severe in high moisture and temperature environments (Czajkowski *et al*. 2009). Infected plants show stunting, chlorosis, wilting, black discoloration and soft rot, which begins at the stem base and progresses upward. Once infection has occurred, the pathogen colonizes the vascular tissues and moves throughout the plant (Raoul des Essarts *et al*. 2016); severe infections lead to whole plant collapse. Latent infections can result in severe losses during storage, especially in warehouses that lack refrigeration facilities (Laurila *et al*. 2008). In recent years, *D. dianthicola* has emerged as one of the most important pathogens infecting potatoes worldwide, and symptoms resemble those of blackleg caused by *Pectobacterium* species (Boluk and Arif 2019). Due to high economic impact in the European potato growing countries, *D. dianthicola* (syn. *Erwinia chrysanthemi* pv. *dianthicola*) has been listed in the EPPO A2 List of pests recommended for regulation as quarantine pests A2/53 (Pg 8); http://archives.eppo.int/EPPOStandards. At present, neither resistant varieties nor effective physical, chemical or biological controls have been reported to manage *D. dianthicola* diseases under field conditions (Czajkowski *et al*. 2014). Only preventive measures are recommended, which include planting certified seeds, eradicating contaminated plant materials, screening for latent infections, monitoring appropriate storage conditions and inspecting fields (Motyka *et al*. 2017). Therefore, early, accurate and sensitive detection of *Dickeya* species is important to identify inoculum sources before they become a problem.

Several assays have been reported for detection of *Dickeya* from infected samples including selective and semi-selective media (Hayman *et al*. 2001; Helias *et al*. 2012), microscopic observations, and biochemical and serological identification methods (Czajkowski *et al*. 2015; Motyka *et al*. 2017). Detection based on molecular methods such as PCR, quantitative PCR and loop-mediated isothermal amplification (LAMP) are implemented routinely due to their high sensitivity, speed, specificity, accuracy, discriminatory ability and reproducibility (Ouyang *et al*. 2013; Czajkowski *et al*. 2015; Ocenar *et al*. 2019). TaqMan probe-based qPCR assays are more accurate, sensitive and reliable for detection of a pathogen (Arif *et al*. 2014). Moreover, addition of 5’ AT-rich flap sequences enhanced the reaction efficiency and sensitivity of multiplex TaqMan qPCR (Arif and Ochoa-Corona 2013; Larrea *et al*. 2019). TaqMan qPCR assays have been reported for the detection of *D. solani* using *fliC* gene, and *D. solani* and *D. dianthicola* using targets based on computational primer prediction pipeline for draft bacterial genomic sequences (Vaerenbergh *et al*. 2012; Pritchard *et al*. 2013). In 2014, van der Wolf *et al*. reported *dnaX* genebased TaqMan qPCR assay for detection of *Dickeya* spp. and described a number of primer pairs to detect different *Dickeya* sp. TaqMan qPCR with the probes labelled with different fluorogenic dyes enables simultaneous detection of multiple pathogens or pests in a single tube (Arif *et al*. 2015). Multiplex TaqMan can be very useful for detection of different *Dickeya* species, which produces nearly indistinguishable disease symptoms without compromising the sensitivity and specificity of the assay.

In this study, we have presented a multiplex TaqMan qPCR for sensitive and simultaneous detection of genus *Dickeya* and *D. dianthicola*. The developed multiplex TaqMan qPCR was validated in the lab extracted DNA from purified cultures and naturally infected potato samples. Comparative genomics using Mauve identified signature genomic regions; the gene *mglA/mglC* region was used to design the primers/probe for the genus *Dickeya*, while the alcohol dehydrogenase gene was used for designing the primers targeting *D. dianthicola*. The developed assay has applications in farm management, seed certification, biosecurity and to discover new reservoir hosts.

## Materials and Methods

### Bacterial strains, growth conditions and plant inoculations in greenhouse

Thirty-three *Dickeya* strains, collected from different geographical locations worldwide, were used in an inclusivity panel (Table 1). Strain numbers prefixed with “A” were obtained from the Pacific Bacterial Collection (University of Hawaii at Manoa). In addition, 34 bacterial strains from other genera comprising closely related species from different genera, were included in an exclusivity panel (Table 1). All strains were stored at −80°C. For culturing, strains were streaked on 2,3,5-triphenyl-tetrazolium chloride sucrose medium (TZC-S: peptone 10 g l^−1^, sucrose 5 g l^−1^, 0.001% TZC and agar 17 g l^−1^) (Norman and Alvarez 1989). The plates were incubated at 26°C (±2°C) overnight; single colonies were streaked on YSC medium (yeast extract 10 g l^−1^, dextrose 20 g l^−1^, calcium carbonate 20 g l^−1^ and agar 17 g l^−1^) and incubated at 26°C (±2°C) overnight; these plates were used for DNA isolation.

**Table 1:**
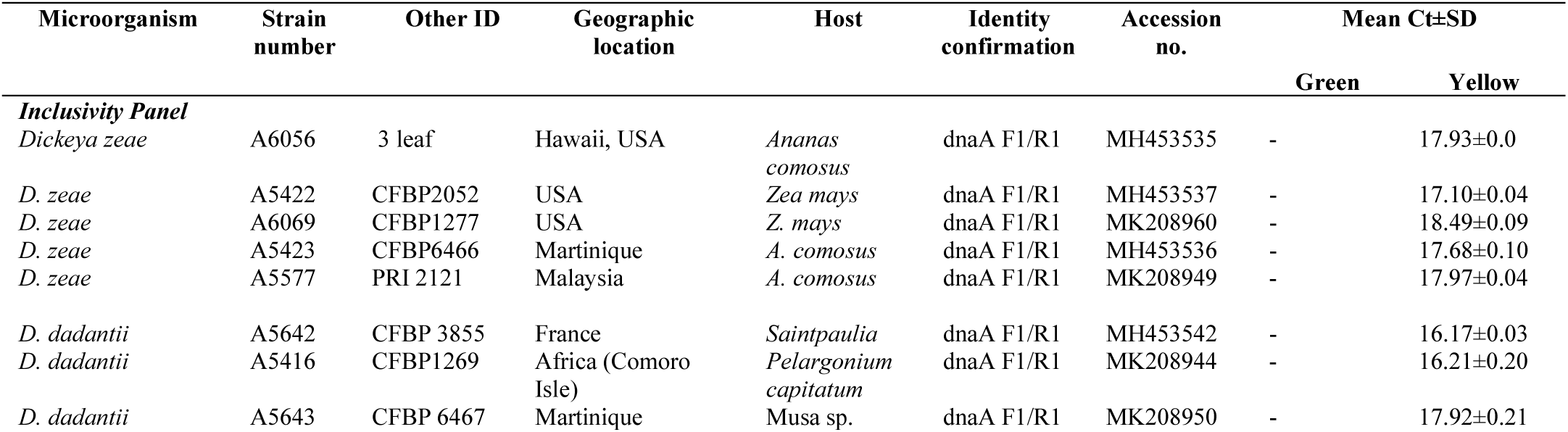
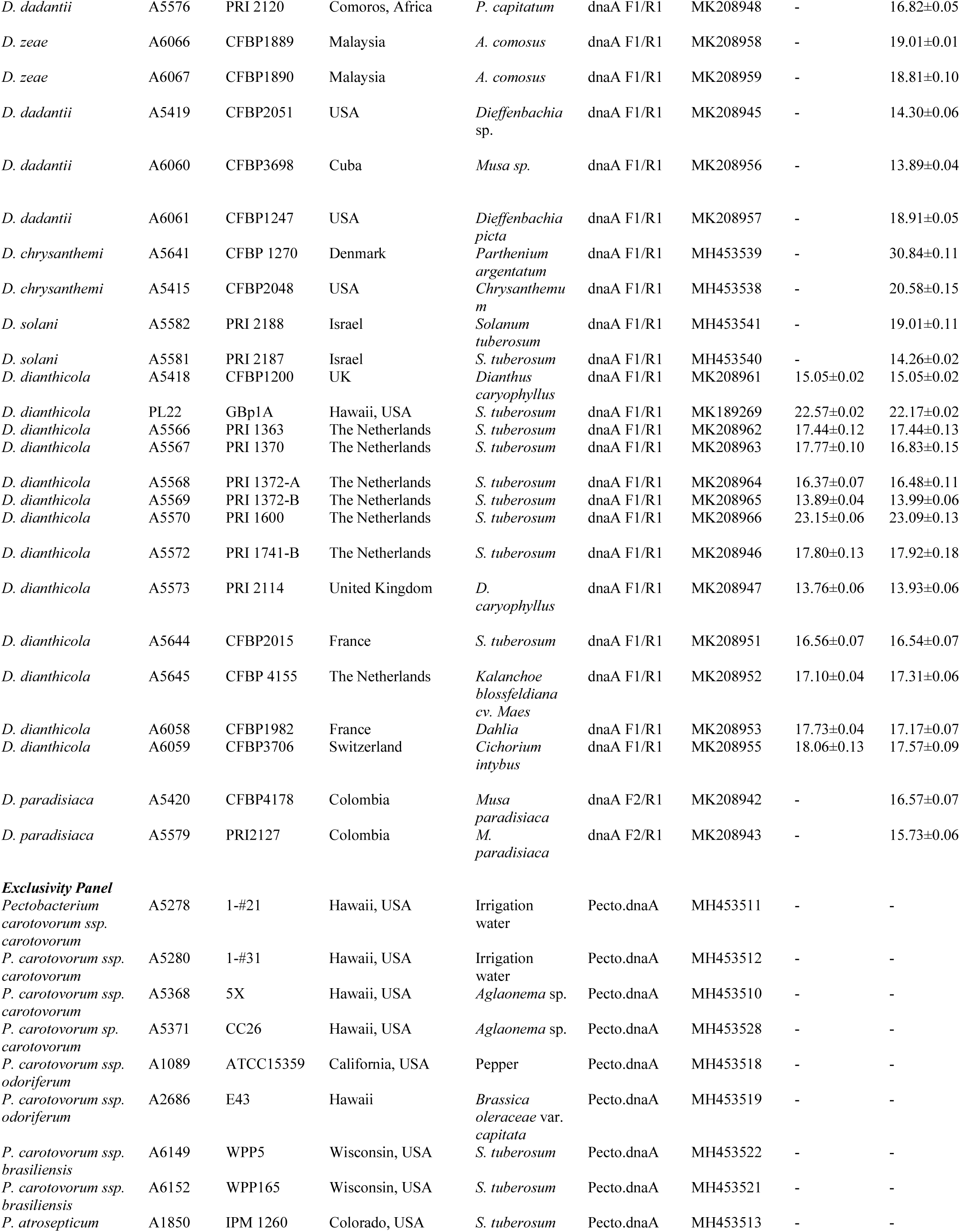
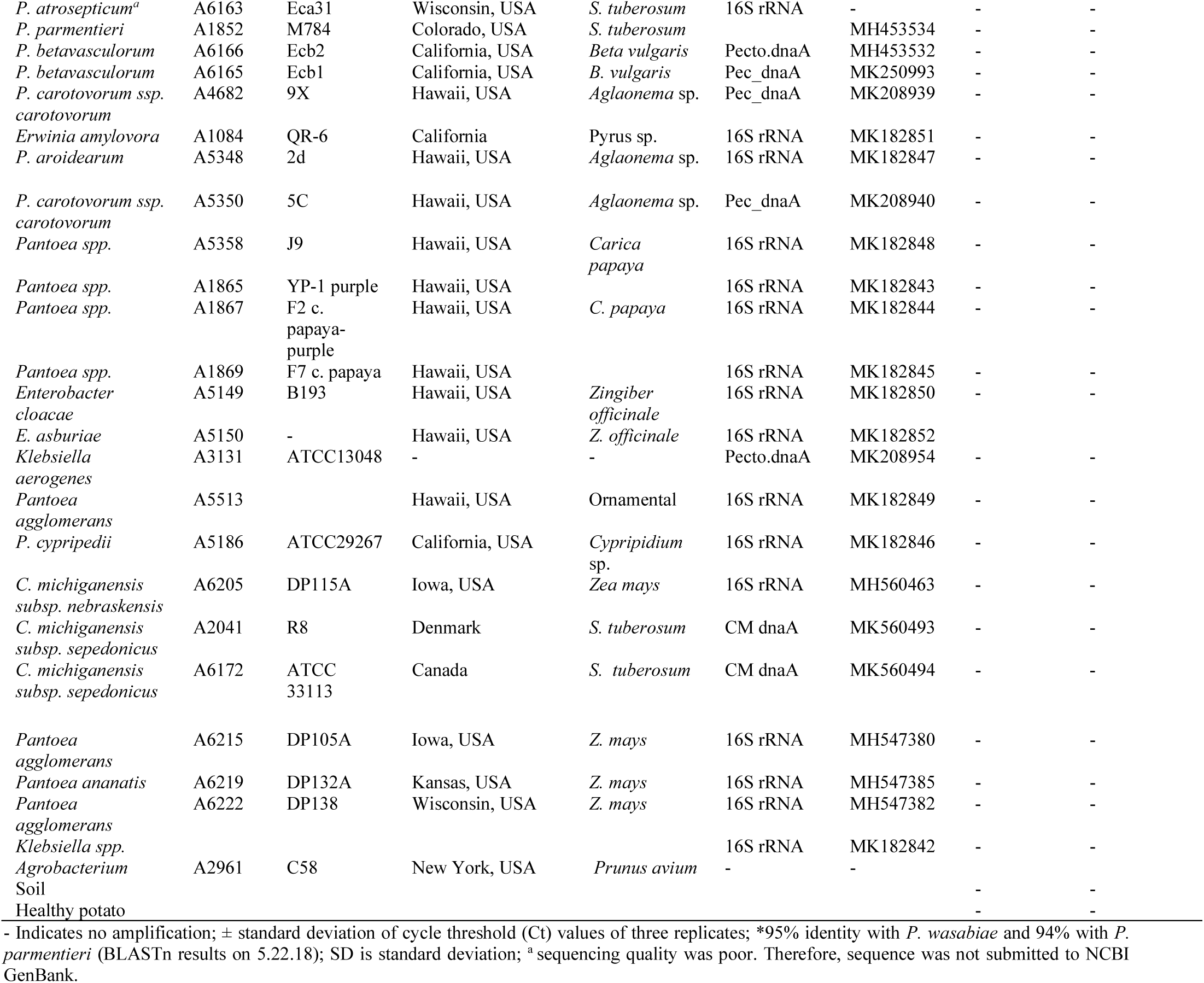
Bacterial strains used for validation of multiplex TaqMan real-time qPCR developed and validated for specific detection of *Dickeya* and *D. dianthicola*.

Healthy potato plants were grown in a greenhouse. Before planting, the sprouting seed potatoes were cut into two pieces, and a piece having at least two buds was planted into a separate pot and placed on the greenhouse bench. Four-week-old healthy potato plants were inoculated with *D. dianthicola* strains. For plant inoculations, sterile scalpels were used to make wounds at the base of the stems of the healthy plants and a loopful of bacteria was stabbed into the fresh wound. The inoculated plants were covered individually with the plastic bags to maintain the humidity, tied and incubated at 28 to 30°C for 24-48 h. Black leg symptoms were observed at the base of the stem three days after inoculation. Healthy plants inoculated with sterile water showed no symptoms. The DNA was extracted from the healthy and inoculated plants.

### Genomic DNA extraction, identification and phylogenetic analysis

Extraction of genomic DNA from infected and healthy plant materials was performed using the Wizard Genomic DNA purification kit (Promega, Madison, WI); DNA was isolated from bacterial cultures using the Ultra Clean Microbial DNA Isolation kit (Mo Bio., Carlsbad, CA). For microbial DNA isolation, 3-4 loopfuls of bacterial cultures from plates were suspended in 300 μl of the microbead solution and vortexed gently to mix; Mini-BeadBeater 16 (Biospec products, Bartlesville, OK) was used to rupture the cell wall at maximum speed for 1 min; bacterial DNA was isolated following the manufacturer’s instructions. DNA concentration and purity were measured using NanoDrop 2000/c spectrophotometers (Thermo Fisher Scientific Inc., Worcester, MA). The genomes of all *Dickeya*, and *Pectobacterium* species were aligned separately with progressive Mauve (Darling et al, 2010); Geneious (version 10.1.3) was used to evaluate the aligned genomes for the *dnaA* (replication initiation factor) regions (data not shown). The primers were designed using Primer3 online software (Rozen, S. & Skaletsky, 2000) following the parameters described by Arif and Ochoa-Corona (2013). The P16s-F1 (5’-AGACTCCTACGGGAGGCAGCA-3’) and P16s-R1 (5’-TTGACGTCATCCCCACCTTCC-3’) were used for 16S region amplification (Dobhal *et al*. 2018). The primer set Dic.dnaA-F1 (5’-TAACAACGTGAACCCCAACGA-3’) and Dic.dnaA-R1 (5’-TCTTCTTTGATGTCGTGACTTTC-3’) targeting *dnaA* region of *Dickeya* was used to amplify all *Dickeya* species except *D. paradisiaca* where Dic.dnaA-F2 (5’-TCCAATGTGAATCCCAAACA-3’) was used in combination with Dic.dnaA-R1. *Pectobacterium* Pec.dnaA-F1 (5’-CATACGTTTGATAACTTCGTTG-3’) and Pec.dnaA-R1 (5’-GATGTCGTGGCTTTCTTCAC-3’) primers were used to amplify the *dnaA* gene region of *Pectobacterium* species. The identity of other bacteria listed in the exclusivity panel was confirmed using primers P16s-F1 and P16s-R1 targeting the 16S ribosomal RNA region. The PCR conditions used for *Dickeya* and *Pectobacterium dnaA* gene amplifications were: Initial denaturation at 95°C for 5 min followed by 35 cycles with 95°C for 20 s, 58°C for 1 min, 72°C for 1 min and a final extension at 72°C for 3 min. The amplified PCR products were cleaned by adding 2 µl ExoSAP-IT (Affymetrix Inc, Santa Clara, CA) in 5 µl of PCR product and incubated at 37°C for 15 min followed by enzyme inactivation at 80°C for 15 min. Sequencing was performed at the GENEWIZ facility (Genewiz, La Jolla, CA) for both sense and anti-sense strands. Obtained sense and anti-sense strands of each strain were aligned, manually edited and the error free consensus sequence was used to confirm identity by comparing the sequences of each strain against the NCBI GenBank nucleotide and genome databases using NCBI BLASTn tool. Sequences generated from each strain were deposited in the NCBI GenBank database and the accessions numbers are listed in Table 1. Consensus sequences of 46 strains used in this study were aligned to generate a phylogenetic tree. Phylogenetic relationships among the strains was generated using the Neighbor-Joining method and the evolutionary distances were calculated using Tamura-Nei method; evolutionary analyses were constructed in MEGA-X (MEGA X; Centre for Evolutionary Functional Genomics) (Saitou and Nei 1987; Felsenstein 1985; Tamura and Nei 1993; Kumar *et al*. 2018). Gaps or undetermined data in the alignment were excluded when site coverage was below 95%. Bootstrap percentages were calculated with 1,000 replicates. The color-coded matrix showing pairwise similarity was generated using Sequence Demarcation Tool v1.2 (Mulshire *et al*. 2014) with the default parameters.

### Target gene selection, genus- and species-specific TaqMan primers and probe design

Genomes of *D. chrysanthemi* strain NCPPB516 (NZ_CM001904), *D. dadantii* strain 3937 (CP002038/NC_014500), *D. dianthicola* strain GBBC 2039 (NZ_CM001838), NCPPB3534 (NZ_CM001840), NCPPB 453 (NZ_CM001841), strain IPO 980 (NZ_CM002023), *D. fangzhongdai* strain DSM 101947 (NZ_CP025003), *D. paradisiaca* strain Ech 703 (CP001654), *D. solani* strain IPO 2222 (NZ_CP015137), strain ND14b (NZ_CP009460), *D. zeae* strain EC1 (NZ_CP006929), strain Ech586 (NC_013592), *Pectobacterium carotovorum* subspecies *carotovorum* strain PCC21 (NZ_018525), *P. atrosepticum* strain 21A (NZ_CP009125), *P. wasabiae* strain CFBP 3304 (NZ_CP015750), *Erwinia amylovora* strain CFBP1430 (NC_013961), *Ralstonia solanacearum* GMI 1000 (NC_003295), and *Clavibacter sepedonicus* strain ATCC33113 (NC_10407) were retrieved from NCBI GenBank Genome database (Supplementary Table 1). Whole genomes were aligned using progressive Mauve (2.4.0) and Geneious (10.2.3). Generated locally Collinear Blocks (LCBs) were analyzed to search unique and conserved regions for the genus *Dickeya* and *D. dianthicola*. The galactose/methyl galactoside ABC transporter ATP-binding protein MglA/MglC region was used to design the primers and probe specific for the genus *Dickeya*. Similarly, for *D. dianthicola*, the genomes of *D. dianthicola* (NZ_CM001838, NZ_CM001840, NZ_CM001841 and NZ_CM002023) were retrieved from the NCBI GenBank Genome database (Supplementary Table 1) and aligned using progressive Mauve; Geneious was used for analyzing the unique genes present in the *D. dianthicola* genome. The alcohol dehydrogenase gene was used to design the primers and probe specific for *D. dianthicola*. The representative genome of each *Dickeya* species and all available complete genomes of *D. dianthicola* along with the genomes of other closely related genera and selected target gene locations were included to generate a BLAST Ring Image Generator (BRIG) 40 (Figure 1) (Alikhan *et al*. 2011). NCBI-BLAST 2.6.0+ database was used to compare and generate the BRIG image. The genus and species-specific primers and fluorescence-labelled probes for the multiplex qPCR assay were designed using Primer3 software and evaluated for thermodynamic characteristics following the parameters described by Arif and Ochoa-Corona (2013). Customized flap sequences were added at 5’ end of each primer designed for genus *Dickeya* and *D. dianthicola* to adjusted Tm, GC content and the length of the primers as previously described (Arif and Ochoa-Corona 2013; Larrea *et al*. 2019). The primers with and without flap sequences along with the thermodynamic parameters are presented in Table 2. No flap sequences were added to the probes. The specificity of each primer was confirmed *in silico* using the BLASTn tool against the NCBI GenBank nucleotide and genome databases (Table 2). All primers and double-quencher probes 5’-/6FAM/ZEN/3IBQ/-3’ and 5’-/5HEX/ZEN/3IBQ/-3’ were synthesized by IDT (Integrated DNA Technologies, Inc., Coralville, IA).

**Table 2:**
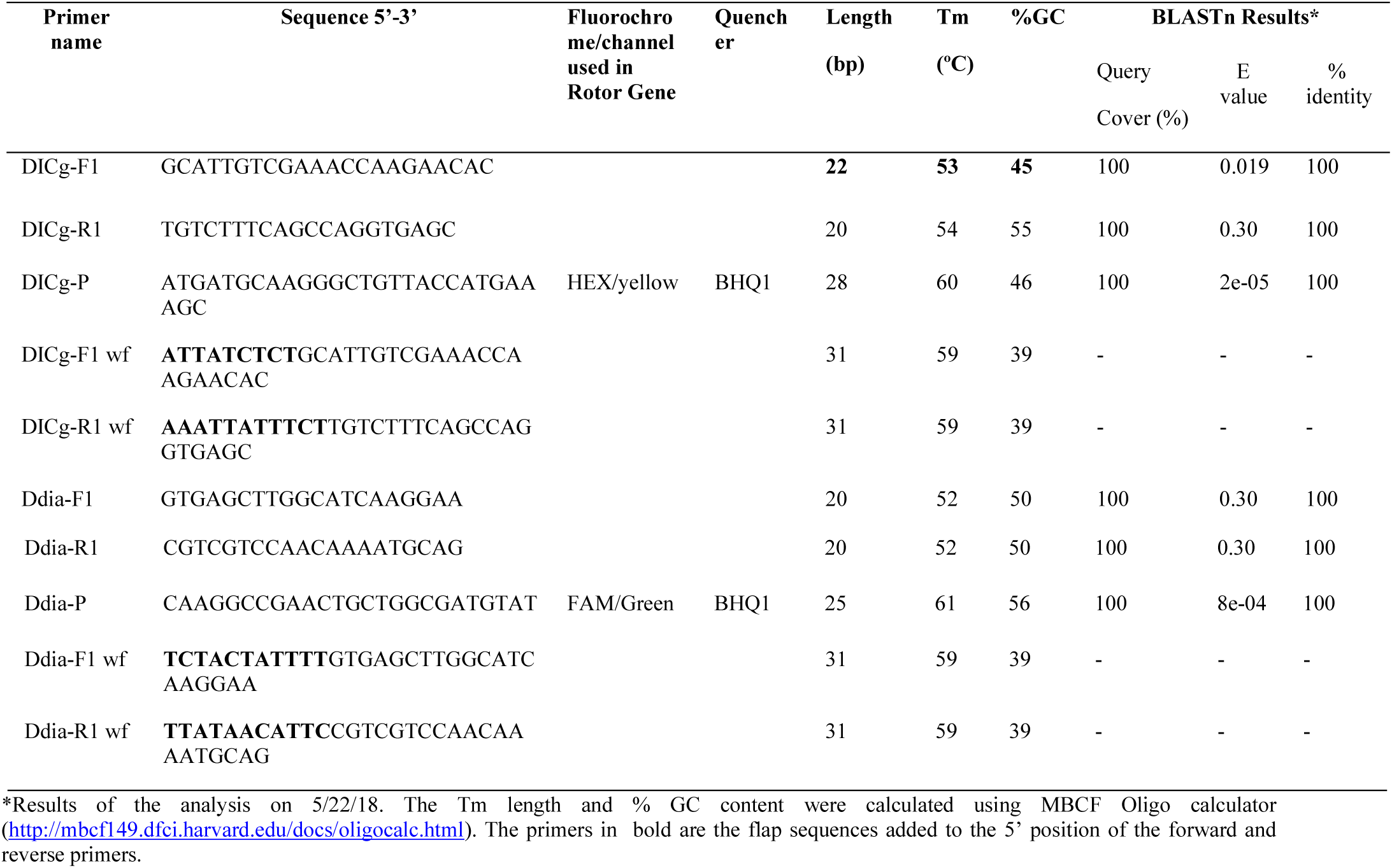
Details of primers and probes used to develop Multiplex TaqMan real-time qPCR assay for specific and rapid detection of all known *Dickeya* species and *D. dianthicola*.

**Figure 1.**
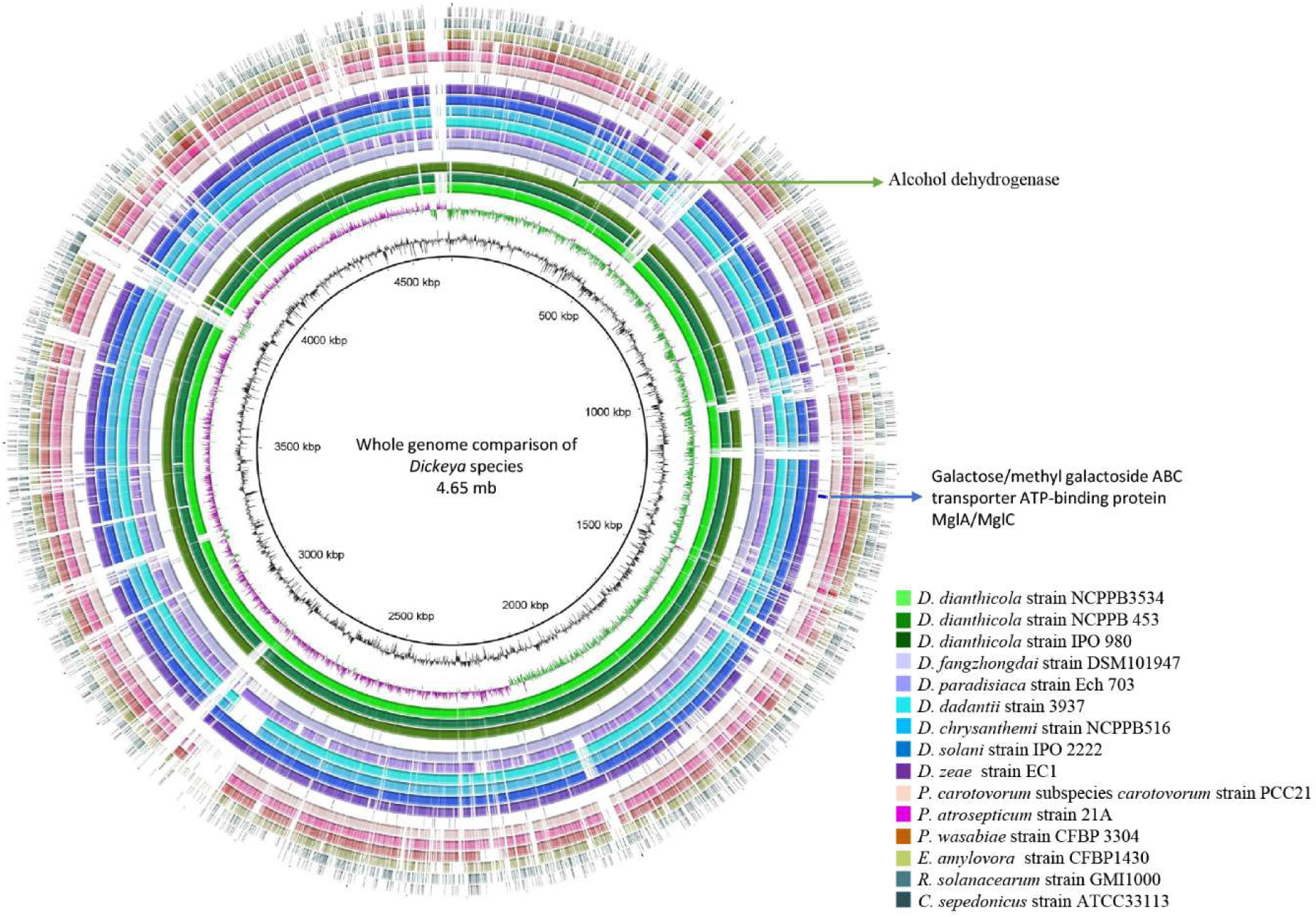
Location of genes used for specific primer and probe design within the whole genome BLAST ring comparison of *Dickeya* species and the closely related bacteria sharing a similar niche with the pathogen. The complete genome of *D. dianthicola* GBBC 2039/LMG25864 (NZ_CM001838) was used as a reference. The GC content and the GC skew (+,-) of the reference genome are represented with the black, purple and green rings. Rings from inside to outside represent the BLAST ring comparison of the whole genomes of *D. dianthicola* strain NCPPB3534 (NZ_CM001840), NCPPB 453 (NZ_CM001841), strain IPO 980 (NZ_CM002023); *D. fangzhongdai* strain DSM 101947 (NZ_CP025003), *D. paradisiaca* strain Ech 703 (NZ_CM001857), *D. dadantii* strain 3937 (CP002038/NC_014500), *D. chrysanthemi* strain NCPPB516 (NZ_CM001904), *D. solani* strain IPO 2222 (NZ_CP015137), *D. zeae* strain EC1 (NZ_CP006929), *Pectobacterium carotovorum* subspecies *carotovorum* strain PCC21 (NZ_018525), *Pectobacterium atrosepticum* strain 21A (NZ_CP009125), *Pectobacterium wasabiae* strain CFBP 3304 (NZ_CP015750), *Erwinia amylovora* strain CFBP1430 (NC_013961), *Ralstonia solanacearum* GMI 1000 (NC_003295), and *Clavibacter sepedonicus* strain ATCC33113 (NC_10407). The blue and green arrows indicate the location of the region for the galactose/methyl galactoside ABC transporter ATP-binding protein MgIA/MglC and alcohol dehydrogenase, respectively. The BLAST RING IMAGE GENERATOR (BRIG) 40 (Alikhan et al., 2011) was used to generate the image. Plasmids were not included in the analysis.

### Multiplex TaqMan real-time qPCR assay

The multiplex TaqMan qPCR was performed in Rotor-Gene Q (Qiagen, Germatown, MD). The concentrations of the primers and probes were adjusted to obtain the optimal amplification. The primer mix was prepared by adding 10 µl of each forward and reverse primer (from stock: 100 µmol l^−1^) for genus *Dickeya* and *D. dianthicola* and 160 µl nuclease free water. The multiplex TaqMan reactions were initially performed in two combinations: primers without 5’ AT-rich flap sequences and with 5’ AT-rich flap sequences. The multiplex TaqMan qPCR reaction was carried out in 25 μl reaction mixture containing 12.5 μl of Rotor-Gene Multiplex PCR Master Mix (Qiagen), 2 μl of the primer mix (5 µmol l^−1^ each), 0.5 μl of each DICg-P and Ddia-P probes (5 µmol l^−1^ stock), 1 µl of template DNA and nuclease free water was adjusted to obtain final volume. Positive and negative controls (non-template; water) were included in each TaqMan qPCR amplification run. Each qPCR reaction was performed in three replicates; standard deviation was calculated. Temperature cycling conditions were: 5 min at 95 °C, followed by 40 cycles of 95 °C for 30 s, and 60 °C for 15 s, acquiring fluorescence on both green (FAM) and yellow (HEX) channels at the end of each extension step. The data analysis was done using the Rotor-Gene Q series software 2.3.1 (Built 49) with auto threshold (Ct); dynamic tube-based normalization was used.

### Exclusivity and inclusivity panels for specificity validation

The specificity of the developed multiplex TaqMan qPCR was validated using the individual DNA template from all 67 bacterial strains included in the inclusivity and exclusivity panels (Table 1). The specificity assays were performed using the primers with flap sequences. All reactions were performed in triplicate; positive and negative template controls were included in each qPCR run. The data analysis was done as described above.

### Multiplex TaqMan qPCR limit of detection determination with and without 5’ AT-rich flap sequences and host background

To minimize the variations among the assays, all sensitivity experiments were performed using the same serial dilutions on the same day. The sensitivity was performed using a ten-fold serially diluted purified genomic DNA (10 ng to 1 fg) of *D. dianthicola* (A5569). To compare the sensitivity, two sets of assays were performed, one using the primer mix with no flap and other using the primer mix with flap primers. The limit of detection of the developed assay was determined in a host DNA background: 1 µl of host (potato) genomic DNA (10 ng µl^−1^) was added in each 10-fold serially diluted genomic DNA of *D. dianthicola* (A5569) from 10 ng to 1 fg. The assays were performed in triplicate. These dilutions along with a no-template control (NTC, water) were run in a Rotor-Gene Q Thermocycler; the TaqMan real-time qPCR conditions were used as described above. The estimated genome size of *D. dianthicola* is ~ 4.86 Mb. Based on the genome size, the copy number was calculated using the formula: number of copies 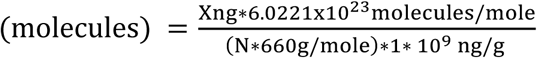 (where X = amount of amplicon (ng), N = length of dsDNA amplicon, 660 g/mole = average mass of 1 bp dsDNA) or web based software (www.scienceprimer.com/copy-number-calculator-for-realtime-pcr).

### Comparison of the sensitivity of the Multiplex TaqMan qPCR with single TaqMan qPCR

To compare the sensitivity of the multiplex with the single TaqMan qPCR assay, the ten-fold serially diluted DNA was used with the single flap primer set. Two separate reaction mixes were prepared, one containing the primer set targeting the genus *Dickeya* and the other targeting *D. dianthicola*. The ten-fold serial dilution sensitivity was performed in the similar way as described above.

### Validation of TaqMan real-time qPCR with naturally infected symptomatic potato samples

The capability of the developed assay was tested with naturally infected potato samples collected from the field. A total of twenty-four plant samples were brought to the lab and washed with sterile water; stem explant of 1 cm size was taken in an Eppendorf tube and crushed with small pestle; DNA was isolated using the Wizard Genomic DNA purification kit. The DNA was quantified using Nanodrop and used for the multiplex TaqMan real-time qPCR. The multiplex TaqMan qPCR was performed using the reaction components and conditions as described above. Three replicates of each sample were run along with the positive and the negative controls.

### Screening of infected plant samples and water sources

The developed TaqMan qPCR was used to screen the samples, collected from six potato growing states of USA, for the genus *Dickeya* and/or *D. dianthicola* contamination/infection. A total of 132 asymptomatic, symptomatic plant samples which included potato stems, tubers and roots, and 3 from potato fields water sources, were screened. The tests were performed at the Wisconsin Seed Potato Certification Laboratory (Wisconsin, USA). Each sample was replicated once; the positive and the negative template controls (NTC) were included in each test run.

## Results

### Genome comparison, primer design and *in-silico* analysis

The highly conserved region present only in the core genome of *D. dianthicola* as well as the genus *Dickeya* was found by evaluating sixteen genomes of the genera *Dickeya, Pectobacerium, Erwinia, Ralstonia* and *Clavibacter* using BLAST comparisons and MAUVE (Figure 1). Mauve-based progressive whole genome alignments enabled the gene selection for the *Dickeya* species and *D. dianthicola*. The alcohol dehydrogenase gene region was identified as a signature gene region present in the core genome of *D. dianthicola* and was used for designing the robust primers and probe specifically amplifying this species. Furthermore, the galactose/methyl galactoside ABC transporter ATP-binding protein mglA/mglC region was identified as a unique target for designing the primers and probe amplifying all known species of genus *Dickeya* (Figure 1). This gene is a part of ABC transporter complex and involved in the import of galactose/methyl galactoside by ATP hydrolysis. Both sets of designed primers and probes were analyzed by BLAST against the NCBI GenBank database and showed 100% query coverage and 100% similarity only corresponding to their target species that is *Dickeya* and *D. dianthicola* (Table 2).

### Identity confirmation and phylogenetic analysis

To determine identities and relationships, all bacterial strains present in inclusivity and exclusivity panels were sequenced. *Pectobacterium* and *Enterobacter* strains were amplified and sequenced using either *dnaA* or 16S rRNA specific primer set to confirm their identity (Table 1). A manually corrected and proofread *dnaA* consensus sequence of ~742 bp was generated. Consensus sequences, produced from both strands, were queried against the NCBI GenBank database using the BLASTn tool to confirm the identity of each strain mentioned in Table 1. The phylogenetic relationships among the bacterial strains were determined using MEGA-X and the sequence demarcation tool (v1.2), clearly grouped the strains of *Dickeya* species from the *Pectobacterium*. All *Dickeya* species strains were clustered together except *D. paradisiaca* which was out-grouped (Figure 2 and 3). The strains of *Pectobacterium* species were clustered all together.

**Figure 2.**
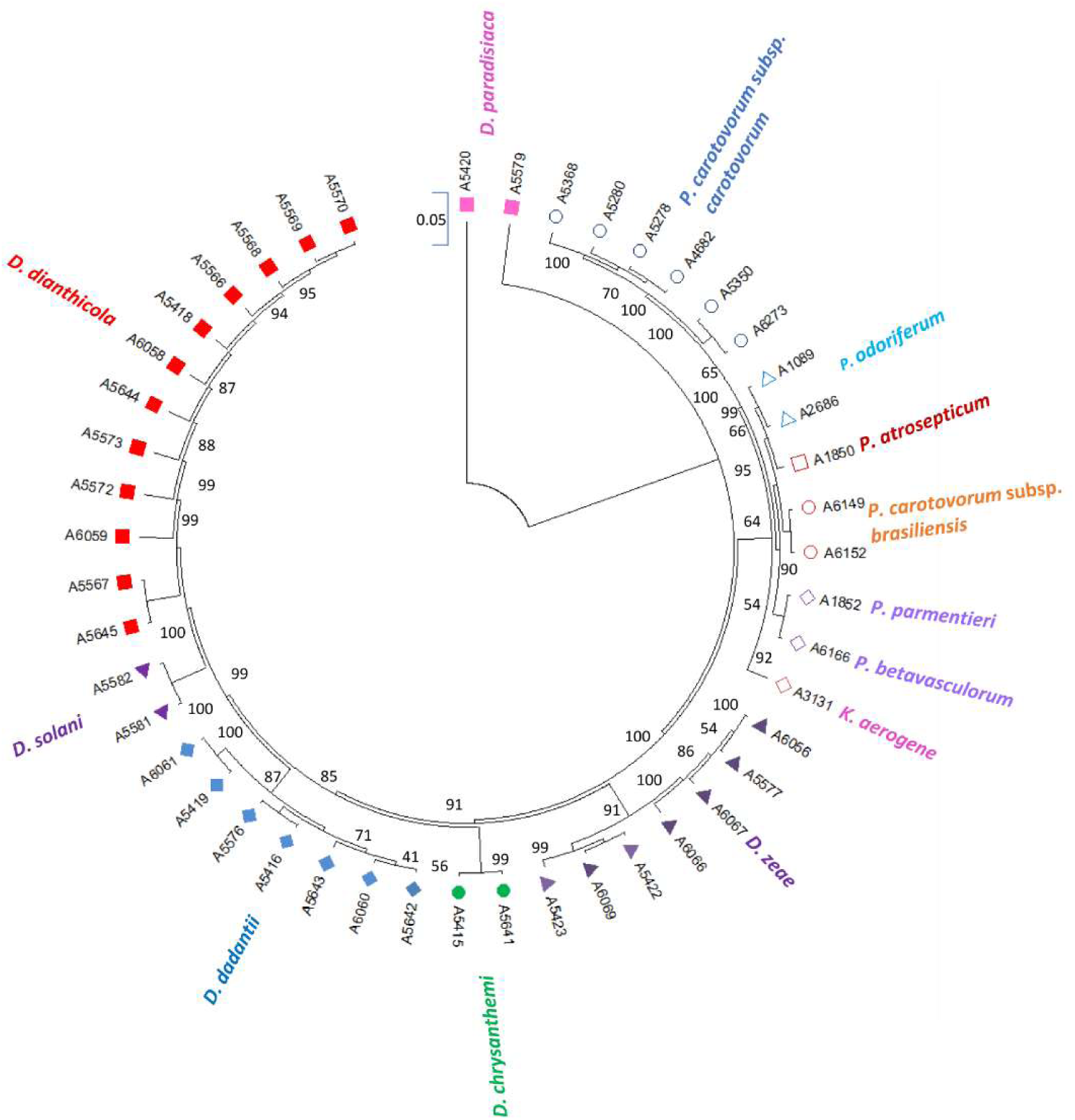
Phylogenetic analyses of *Dickeya* strains using *dnaA* gene conducted in MEGA X. The analysis was inferred using the Neighbor-Joining method. The optimal tree with the sum of branch length = 0.91379220 is shown. The percentage of replicate trees in which the associated taxa clustered together in the bootstrap test (1000 replicates) are shown next to the branches. The evolutionary distances were computed using the Tamura-Nei method and are in the units of the number of base substitutions per site. Evolutionary analyses were conducted in MEGA X. The analysis involved 46 nucleotide sequences which includes *Dickeya, Pectobacterium* and *Enterobacter* species. All *Dickeya* strains grouped together except two *D. paradisiaca* strains that form a separate clade. All *Pectobacterium* species clustered together.

**Figure 3.**
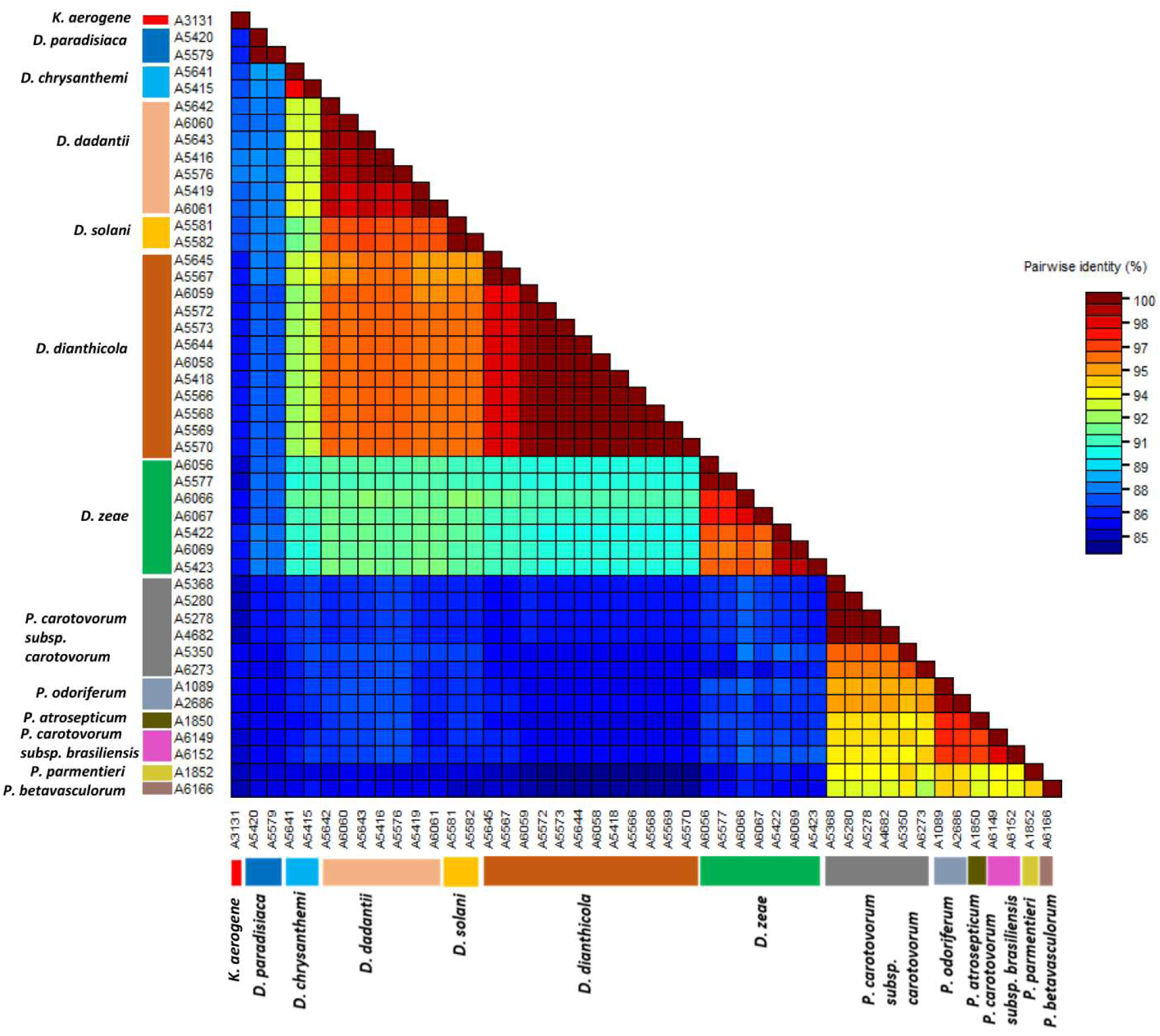
Color-coded matrix showing pairwise identity scores between *dnaA* gene sequences of strains of *Dickeya* sp. with strains of other *Pectobacterium* sp. and *Enterobacter aerogenes*. species. Forty-six partial sequences were used for constructing the pairwise similarity scores displaying a color-coded matrix considering the number of mismatching and the number of positions along the alignment.

### Specificity of the TaqMan primers and probes

The *in-silico* analyses were performed via similarity search of each primer sequence against the NCBI GenBank database using the BLASTn tool; primers showed 100% specificity with query coverage and percent identity match of 100% with the target *Dickeya* genus and the species *dianthicola* (Table 2). During *in-vitro* experiments, the specificity of the developed assay was tested with the DNA from 13 and 20 strains of *D. dianthicola* and other species of *Dickeya* from distinct geographical origins, respectively (Table 1). The Ct values from *Dickeya* strains, excluding *D. dianthicola*, ranged from 13.89±0.04 to 30.84±0.11 in the yellow channel (n=18), while the green channel showed amplification and Ct values from 13.76±0.06 to 23.15±0.06 with the strains of *D. dianthicola* (n=13). The yellow channel exhibited almost similar Ct values as the green channel for the same strains of *D. dianthicola* (13.93±0.06 to 23.09±0.13). No amplification in green or yellow channels were observed from the non-target and/or closely related species (n=34). This demonstrated that the developed primers/probes are highly specific for the genus *Dickeya* and specifically for *D. dianthicola* (Table 1). No Ct values were observed in either green or yellow channels with any member of the exclusivity panel, healthy plant or soil sample included in the assay validation. In addition, during the specificity validation, no Ct value >35 was obtained with any member of the inclusivity panel. The amplification from each representative species from genus *Dickeya* is shown in Figure 4. The primers with and without 5’ AT-rich flap sequences were also tested for their specificity; no false positives or false negatives were obtained (data not shown). The assays were highly specific but still the samples with Ct value after 35 cycles were considered negative.

**Figure 4.**
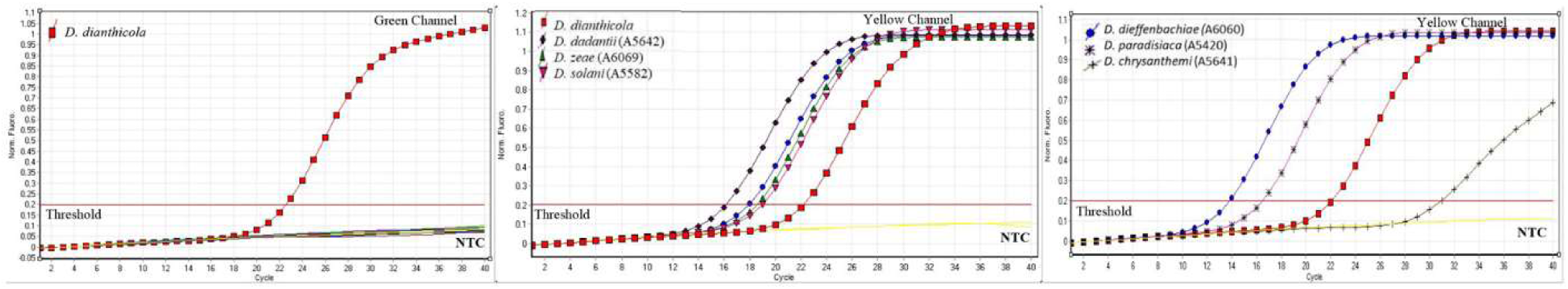
Multiplex TaqMan qPCR assay showing representative strains of *Dickeya* species included in the inclusivity panel. The green and yellow channels correspond to the different reporter dyes 6-FAM (495/520) and HEX (535/554) (excitation/emission spectra in nm) for detection of genus and species of *Dickeya*, respectively. NTC, Non-template control. Each reaction was performed in triplicate.

### Limit of detection determination

The primer sets with and without flap sequences were compared for their sensitivity using the purified *D. dianthicola* genomic DNA. Both primer sets with and without flaps were sensitive and detected 10 fg of purified genomic DNA (Figure 5A1, A2; 5B1, B2). The primers with the 5’ AT-rich flap showed better reaction efficiency compared to the primer set with no flap (Figure 5A1, A2). At lower concentrations of template DNA, the primers with no AT-rich flap sequences showed higher standard deviation among the replicates compared to primers with flap, indicating the enhanced stability and better reaction thermodynamics.

**Figure 5.**
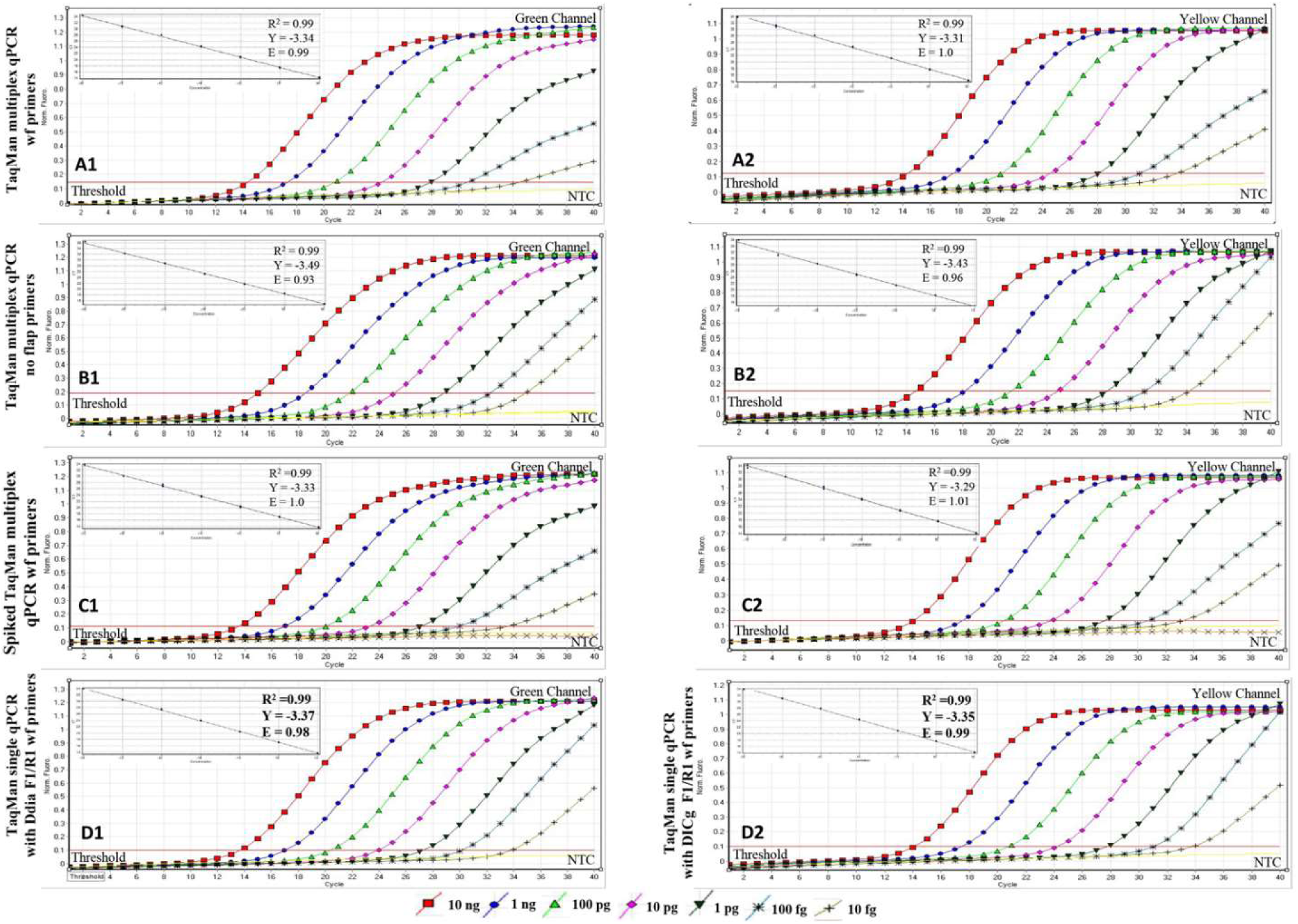
Multiplex and single TaqMan qPCR standard curves and graphs. The graphs and curves were generated using ten-fold serial diluted genomic DNA (10 ng to 1 fg) of *Dickeya dianthicola* (A, B, D) and ten-fold serially diluted genomic DNA mixed with host plant DNA (C). The Green and yellow channels correspond to the different reporter dye 6-FAM (495/520) and HEX (535/554) (excitation/emission spectra in nm), respectively. A1/A2 multiplex TaqMan qPCR using DICg-F1/R1wf and Ddia F1/R1 wf (wf -with flap primers) primer sets in the reaction mix. B1/B2 multiplex TaqMan qPCR using DICg-F1/R1 and Ddia F1/R1 (no flap primers) primer sets in the reaction mix. C1/C2 spiked multiplex TaqMan qPCR using DICg-F1/R1wf and Ddia F1/R1 wf (wf -with flap primers) primer sets in the reaction mix (the spiked assay was done by adding 1 μl of healthy plant-potato DNA extracted from tubers to each serial dilution to simulate natural infection and then performing the multiplex qPCR assay). D1 single TaqMan qPCR with Ddia F1/R1 wf primer set in a reaction mix. D2 single TaqMan qPCR with DICg F1/R1 wf primer set in a reaction mix. The C_T_ values represent the average of three replicates ±SD. Slopes (Y = threshold cycles (Ct) of target DNA detected), R^2^ (correlation coefficient), and E (= amplification efficiency of the real-time PCR assay) of each reaction are presented in the respective curves. X axis represents the number of cycles, and Y axis is normalized fluorescence.

No change in the sensitivity was observed when 1 µl of host genomic DNA (DNA extracted from healthy potato plant) was added to each ten-fold serially diluted genomic DNA. The detection limit of 10 fg was detected; demonstrating the robustness of the developed assay with no adverse effect from host background DNA (Figure 5C1, C2).

To demonstrate the robustness of the multiplex TaqMan qPCR, a single TaqMan qPCR assay was also performed to compare the detection limit using both primer/probe sets. Primer sets DICg-F1/R1-wf and Ddia-F1/R1-wf were able to detect up to 10 fg in both yellow and green reporting channels, respectively. These results indicated that the developed multiplex assay is robust and equally sensitive as single target TaqMan real-time qPCR assay (Figure 5D1, D2).

### Multiplex TaqMan qPCR validation with naturally infected plant samples

A total of 13 naturally infected asymptomatic and symptomatic plants were tested positive using the developed multiplex TaqMan real-time qPCR; all samples gave positive results using both the *Dickeya* genus and *D. dianthicola* specific primer sets. The Ct values in both yellow and green channels ranged from 18.02±0.04 to 32.37±0.52 and 18.16±0.01 to 34.08±1.07, respectively. The bacteria were isolated from the positive samples and sequenced with forward and reverse primers described previously to confirm their identity. The multiplex qPCR showed 100% accuracy in detection of the *Dickeya* infected field samples. No false positives or false negatives were detected during the experiment (Figure 6).

**Figure 6.**
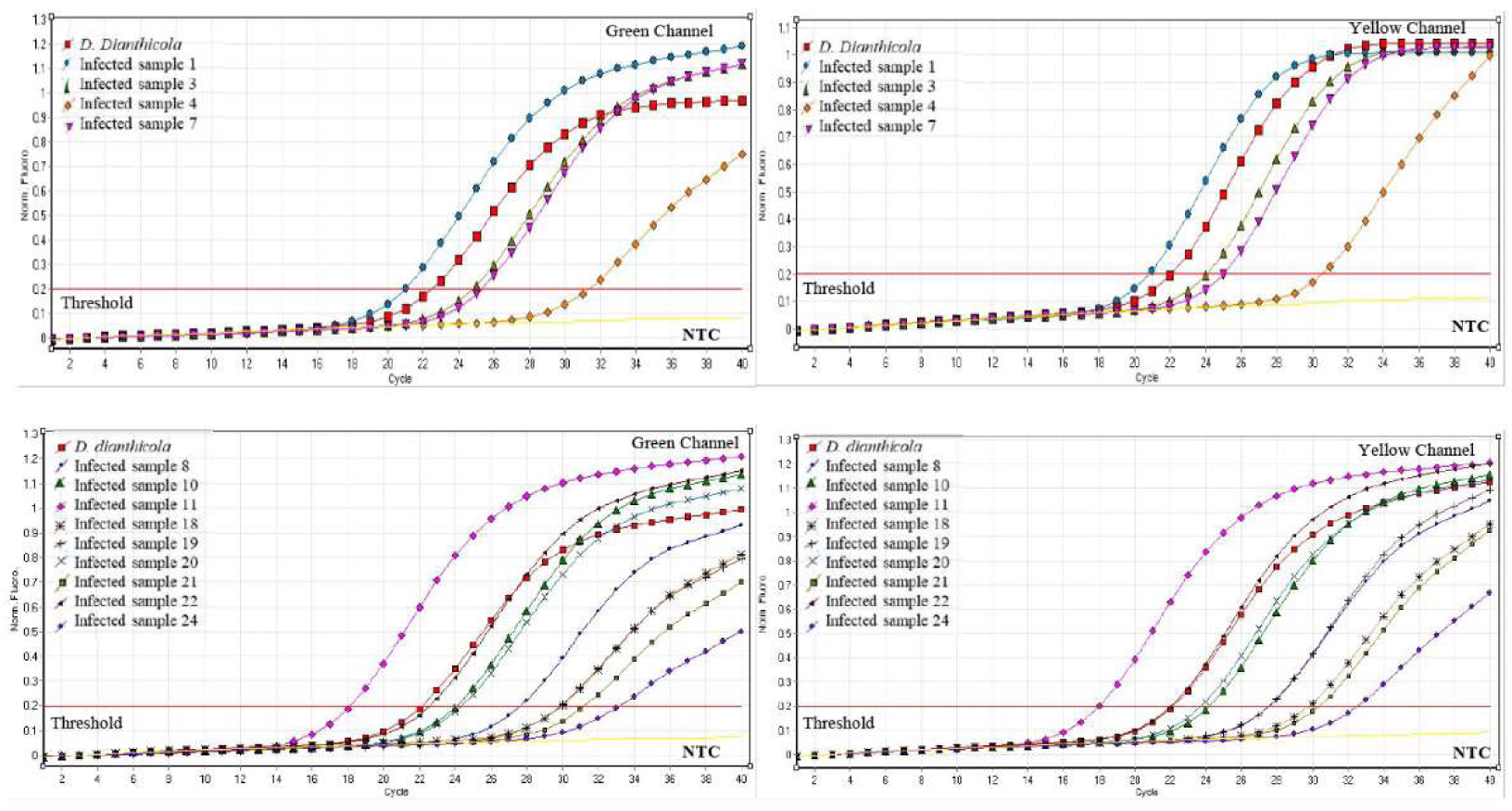
Multiplex TaqMan qPCR assay using infected field samples. The Green and yellow channels correspond to the different reporter dye 6-FAM (495/520) and HEX (535/554) (excitation/emission spectra in nm), respectively. *D. dianthicola*, Positive control; Infected potato plant samples 1, 3, 4, 7, 8, 10, 11, 18, 19, 20, 21, 22 and 24; NTC, Non-template control. Each reaction was performed in three replicates.

### Multiplex TaqMan qPCR assay validation with different sample types collected from multiple locations

The developed multiplex TaqMan real-time qPCR was used to screen 143 field samples (stem, potato tuber, root and water) for *Dickeya* and specifically for *D. dianthicola* infection (Table 3). Out of 143 samples, 61 were positive for genus *Dickeya* and of these, 60 were positive for *D. dianthicola*. Samples crossing the threshold value (Ct) after 35 cycles were considered negative.

**Table 3:** Samples tested during performance evaluation of the multiplex TaqMan real-time qPCR for specific detection of *Dickeya* species and *D. dianthicola* from naturally infected potato tissues.

| Sample Type | Location/State | Total number of samples tested positive/total number of samples tested (%) |  |
| --- | --- | --- | --- |
|  |  | DICg-F1/R1 wf<br>(*Dickeya* genus) | Ddia-F1/R1 wf<br>(*D. dianthicola*) |
| *Plant samples*<br>(includes plant parts) |  |  |  |
| Stems | Missouri | 19/20 (95%) | 19/20 (95%) |
| Stems | North Carolina | 5/5 (100%) | 5/5 (100%) |
| Stems | Indiana | 3/3 (100%) | 3/3 (100%) |
| Stems | Florida | 14/14 (100%) | 14/14 (100%) |
| Stems | Texas | 3/3 (100%) | 3/3 (100%) |
| Stems | Idaho | 3/3 (100%) | 3/3 (100%) |
| Stems | Wisconsin | 3/21 (14%) | 3/21 (14%) |
| Roots | Wisconsin | 2/2 (100%) | 2/2 (100%) |
| Tuber | Wisconsin | 18/56 (31%) | 18/56 (31%) |
| Tuber | Florida | 2/2 (100%) | 2/2 (100%) |
| *Other Sources* |  |  |  |
| Water | Wisconsin | 1/3 | 2/3 |
| FTA cards | Wisconsin | 5/11 (45%) | 3/11 (27%) |

## Discussion

In plant disease management, rapid, sensitive and accurate detection of pathogens is critical for timely responses that facilitate disease control. Furthermore, early disease detection is important for assessing plant health prior to seedling transplant. In this study, we developed a robust multiplex TaqMan real-time qPCR for simultaneous detection of all known *Dickeya* species and *D. dianthicola* in a single reaction. The primers were developed from unique and conserved genes, extracted using comparative genomic analyses of *Dickeya* species. The developed method has been thoroughly validated using strains in inclusivity and exclusivity panels as well as naturally and artificially infected samples. The method showed 100% accuracy and successfully detected infection of both *Dickeya* species and *D. dianthicola* from infected plant samples received from different states of the United States.

*Dickeya* is one of the most important enteric soft rot pathogens known to cause tremendous economic losses worldwide. *Dickeya* species, especially *D. dianthicola*, is a major problem for potato growing countries, and therefore is included in the European Union A2 quarantine list of pathogens. Accurate detection, distribution and monitoring of this pathogen is critical for prevention of higher economic losses. Detection by culturing and isolation is time-consuming, especially when the pathogen can be overgrown by saprophytic bacteria commonly present in infected samples. PCR provides fast, robust and accurate detection of pathogens compared to traditional culture-based methods. Compared to traditional PCR, real-time qPCR allows faster, accurate, sensitive and quantitative monitoring of a pathogen with less chance of carry over contaminations (Arif *et al*. 2014, 2015; Strayer *et al*. 2016; Larrea *et al*. 2019). TaqMan real-time qPCRs have been developed for the detection of *D. solani* and *D. dianthicola* (Van Vaerenbergh *et al*. 2012; Pritchard *et al*. 2013), but these protocols have not been thoroughly validated. TaqMan qPCR assays for *Dickeya* species using the *dnaX* gene region have been reported by van der Wolf *et al*. (2014). However, the assay developed using *dnaX* gene sequence for detection of *D. solani* was insufficiently specific and found not to be suitable for testing propagative material (van der Wolf *et al*. 2014). Therefore, the whole genome comparative genomics approach was used to develop robust primers and probe sets using highly unique and conserved genes. Multiplexing the reaction using different probes labeled with different fluorogenic dyes allows detection of multiple targets in a single reaction which facilitates rapid detection, reduces assay costs and enhances assay reliability (Arif *et al*. 2015; Larrea *et al*. 2019). The sensitivity and specificity of the developed assays depends on the target gene selection and the thermodynamics of the primers and probes (Arif and Ochoa Corona 2013).

During this study, comparative genomics identified unique region/gene in the target genomes of *Dickeya* species. We designed the primers from these target genes and their flanked region – this approach enhances the specificity and reliability of the developed assay since the gene arrangement among the strains within a species is conserved compared to strains from different species. The primers were selectively tested both *in-silico*, by searching for the homology in the nucleotide database (NCBI GenBank BLASTn), and *in-vitro*, by screening against the strains present in inclusivity and exclusivity panels (Table 1). Using BLASTn, the primers and probes were found to be specific for the target pathogen with 100% identity match and query coverage. No false negatives or false positives were observed during the validation. In our lab, the alcohol dehydrogenase gene was also used to develop a loop-mediated isothermal amplification assay for specific, on-site detection of *D. dianthicola* and proved to be highly specific (Ocenar *et al*. 2019).

Previously, we have demonstrated that incorporation of flap sequences at the 5’ position of the primer could lead to increased sensitivity, fluorescence, overall PCR yield and optimize reaction efficiency (Arif and Ochoa-Corona 2013; Larrea *et al*. 2019). The 5’ AT-rich flap sequences were added to both primers sets, DICg F1/R1 and Ddia F1/R1, to evaluate their effects on sensitivity and/or specificity of the developed assay. Both primers with and without 5’ AT-rich flap sequences, targeting the *Dickeya* species and *D. dianthicola*, were specific for respective pathogens; no cross-reactions were observed (data not shown). Both multiplex primer sets, DICg F1/R1wf and Ddia F1/R1wf, with 5’ AT-rich flap showed optimum reaction efficiency compared to the primers with no flaps (Figure 5) and were sufficiently sensitive to detect the genomic DNA of *D. dianthicola* and *Dickeya* species down to 10 fg. However, primers with no flap showed some discrepancy among the replicates at the lowest concentration (10 fg; DICg F1/R1). Therefore, both primers with the 5’ flap sequences were used for overall validation of the developed multiplex TaqMan qPCR assay. The addition of the 5’ AT-rich flap also made the primers and probes compatible to mix and match based on user’s need without losing specificity and/or sensitivity; these primers work under the “One Lab - One Protocol” concept (Arif 2019).

Multiplexing for specific detection of multiple targets in a single reaction is time efficient, provides enhanced reliability and further validation of the outcomes (Li *et al*. 2017). The probes labelled with internal ZEN quencher along with the traditional 3’end quencher enhance quenching efficiency, provide less background noise and increase precision (Arif *et al*. 2013). Multiplexing the reaction with different primer and probe sets can adversely affect the sensitivity and can increase the background noise (Li *et al*. 2017). However, the developed assay did not show any discrepancy in the detection limit when assays were performed individually as compared to the multiplex reaction; both detected 10 fg of genomic DNA. The presence of plant DNA mixed with pathogen DNA has been reported to interfere with the *Taq* DNA polymerase due to the presence of strong DNA-binding domains, reducing *Taq-*polymerase availability and activity, which results in higher Ct values (Hoy *et al*. 2001; Strayer *et al*. 2016). Therefore, the spiked assays were performed by adding the DNA from healthy plants along with target pathogen DNA in the multiplex qPCR reaction. No adverse effect on the Ct values was observed – the detection limit remained the same (10 fg) as with the pure genomic DNA from *D. dianthicola*. The genome size of *Dickeya* species ranges from 4.62-5.05 Mb (supplementary Table 1). Based on the calculations, 10 ng of *D. dianthicola* genomic DNA contains 1.93×10^6^ copies, therefore, 10 fg should be equivalent to ~ 2 copies. The sensitivity of the developed multiplex TaqMan qPCR is higher (10 fg per reaction; Ct=33.75±0.24 compared to the previously reported primers/assays by Potrykus *et al*. (2014) for *Dickeya* species (detection limit 10 pg per microlitre) and van der Wolf et al. (2014) for *D. dianthicola* (20 fg per reaction; Ct=36.5±0.38).

The effectiveness and robustness of any developed method is accessed by its ability to detect the pathogen from naturally infected samples. Therefore, the developed assay was tested with twentyfour asymptomatic and symptomatic, naturally infected plant samples. Thirteen samples showed positive amplification in both yellow and green channels, confirming the presence of *D. dianthicola*. Each infected sample that tested positive with TaqMan qPCR assay was further confirmed by sequencing the *dnaA* gene region followed by BLASTn (data not included). The BLASTn results confirmed 100% identity match with *D. dianthicola*. Furthermore, the developed method was also tested with 143 different plant and water samples received from various locations by the Wisconsin Potato certification laboratory. The method was able to detect 61 samples positive for *Dickeya* species and 60 for *D. dianthicola*. The developed method demonstrated the ability to be used for monitoring *Dickeya* species and *D. dianthicola* infections in naturally infected plant samples.

In conclusion, we developed a highly sensitive, robust, accurate, reproducible and specific multiplex TaqMan qPCR based on the whole genome comparative genomics approach, to simultaneously detect *Dickeya* species and *D. dianthicola*. To our knowledge, this is the only multiplex TaqMan qPCR to simultaneously detect genus *Dickeya* and *D. dianthicola*. The potato industry involves the global exchange of plant propagative materials; this tool will allow diagnosticians and certification agencies/labs to quickly test and identify the pathogen from pure culture and/or infected plant samples to prevent the dissemination of the disease to uninfected areas. Practical applications of the developed assay were demonstrated in rapid and large-scale screening of potato samples for certification purposes. This tool can also be utilized for the phytosanitary, disease epidemiology and management purposes.

## Acknowledgments

This work was supported by the USDA National Institute of Food and Agriculture, Hatch project 9038H, managed by the College of Tropical Agriculture and Human Resources, and USDA FARMBILL (project number: APP-7058). The strains used in this study were revived and characterized with grant support from the National Science Foundation (NSF-CSBR grant no. DBI-1561663). The mention of trade names or commercial products in this publication does not imply recommendation or endorsement by the University of Hawaii. The findings and conclusions in this publication are those of the authors and should not be construed to represent any official USDA or U.S. Government determination or policy. This research was supported in part by the U.S. Department of Agriculture, Animal and Plant Health Inspection Service.

## Supplementary Materials

**Supplementary Table 1**. Details of genomes used for target gene selection for development of TaqMan qPCR for specific detection of the Genus *Dickeya*.

