## Supplementary Table 1 for "Comparative genomics reveals signature regions used to develop a robust and sensitive multiplex TaqMan real-time qPCR assay to detect the genus *Dickeya* and *Dickeya dianthicola*"

**Supplementary Table 1.** Details of genomes used for target gene selection for development of TaqMan qPCR for specific detection of the Genus *Dickeya*.

| Organism | GenBank Accession Number | Genome Size (mb) | No. of Plasmids/ unknown | GC % | Strain I.D. | Host | Location | Sequencing Platform/Institute |
|---|---|---|---|---|---|---|---|---|
| *Dickeya chrysanthemi* | NZ_CM001904 | 4.62 | 1 | 54.24 | NCPPB 516 | *Parthenium argentatum* | Denmark | 454 |
| *D. dadantii* | CP002038/NC_014500 | 4.92 | - | 56.30 | 3937 | *Saintpaulia ionantha* | USA | |
| *D. dianthicola* | NZ_CM001838 | 4.79 | 1 | 56.26 | GBBC 2039/LMG25864 | *Solanum tuberosum* 'Première' | Belgium | Illumina |
| *D. dianthicola* | NZ_CM001840 | 4.86 | 1 | 55.64 | NCPPB 3534 | *S. tuberosum* | the Netherlands | 454; Illumina |
| *D. dianthicola* | NZ_CM001841 | 4.67 | 1 | 55.95 | NCPPB 453 | *Dianthus caryophyllus* | UK | 454 |
| *D. dianthicola* | NZ_CM002023 | 4.84 | 1 | 55.70 | IPO 980 | *S. tuberosum* | Netherlands | 454 |
| *D. paradisiaca* | CP001654 | 4.67 | - | 55.00 | Ech703 | *S. tuberosum* | USA | - |
| *D. solani* | NZ_CP009460 | 5.05 | - | | ND14b | waterfall | Malaysia | PacBio RSII |
| *D. solani* | NZ_CP015137 | 4.92 | - | 56.20 | IPO 2222 | *S. tuberosum* | Netherlands | Illumina NextSeq 500; PacBio |
| *D. zeae* | NZ_CP006929 | 4.53 | - | 53.40 | EC1 | Rice | China | Illumina HiSeq |
| *D. zeae* | NC_013592 | 4.81 | - | 53.60 | Ech586 | | USA | |
| *P. carotovorum* subsp. *carotovorum* | NZ_018525 | 4.84 | - | 52.20 | PCC21 | Chinese Cabbage | Korea | Roche 454 GS FLX |
| *P. atrosepticum* | NZ_CP009125 | 5.02 | 1 | 51.10 | 21A | *S. tuberosum* | Belarus | Illumina MiSeq |
| *P. wasabiae* | NZ_CP015750 | 5.04 | - | 50.60 | CFBP 3304 | *Eutrema wasabi* | Japan | PacBio |
| *E. amylovora* | NC_013961 | 3.83 | 1 | 53.57 | CFBP1430 | *Crataegus oxyacantha* | France | Illumina |
| *R. solanacearum* | NC_003295 | 5.81 | 1 | 66.96 | GMI1000 | *S. lycopersicum* | France | - |
| *Clavibacter sepedonicus* | NC_10407 | 3.40 | 2 | 72.42 | ATCC33113 | *S. tuberosum* | Canada | - |
